# µPhos: a scalable and sensitive platform for functional phosphoproteomics

**DOI:** 10.1101/2023.04.04.535617

**Authors:** Denys Oliinyk, Andreas Will, Felix R. Schneidmadel, Sean J. Humphrey, Florian Meier

**Author notes:** correspondence to: Prof. Dr. Florian Meier (Jun.-Prof.), Dr. Sean J. Humphrey.

## Abstract

Mass spectrometry has revolutionized cell signaling research by vastly simplifying the identification and quantification of many thousands of phosphorylation sites in the human proteome. Defining the cellular response to internal or external perturbations in space and time is crucial for further illuminating functionality of the phosphoproteome. Here we describe µPhos, an accessible phosphoproteomics platform that permits phosphopeptide enrichment from 96-well cell culture experiments in < 8 hours total processing time. By minimizing transfer steps and reducing liquid volumes to < 200 µL, we demonstrate increased sensitivity, over 90% selectivity, and excellent quantitative reproducibility. Employing highly sensitive trapped ion mobility mass spectrometry, we quantify more than 20,000 unique phosphopeptides in a human cancer cell line using 20 µg starting material, and confidently localize > 5,000 phosphorylation sites from 5 µg. This depth covers key intracellular signaling pathways, rendering sample-limited applications and extensive perturbation experiments with hundreds of samples viable.

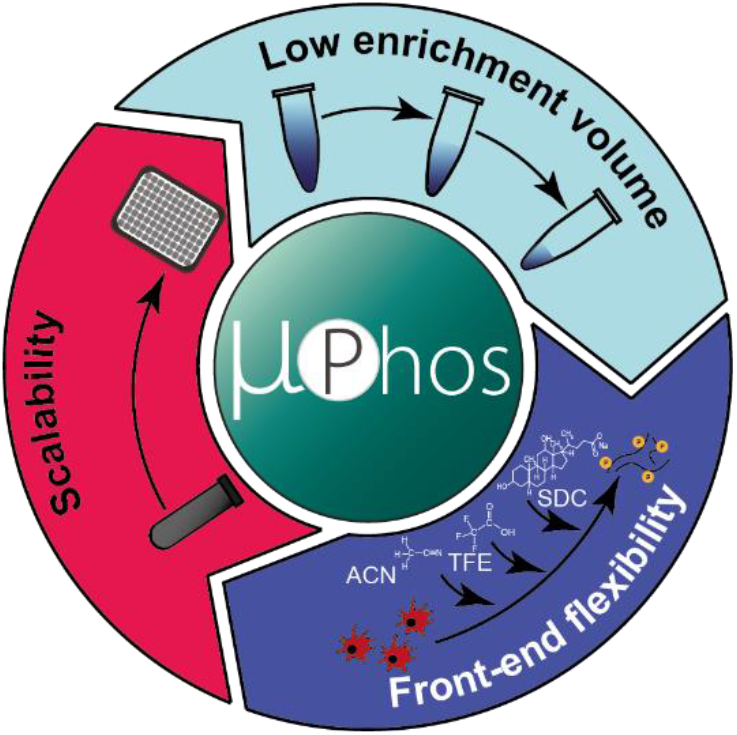

## Introduction

Protein phosphorylation is a widespread post-translational modification (PTM) that reversibly regulates cellular processes through a complex intracellular network of kinases and phosphatases^1^. At least three quarters of proteins in the cell are phosphorylated and dysregulated phosphorylation is associated with numerous complex diseases including cancer^2–7^. Advances in mass spectrometry (MS) instrumentation, sample preparation and bioinformatics have enabled the study of protein phosphorylation dynamics on a system-wide scale^8,9^. As witnessed in proteomics, where rapid high-coverage proteomes can be obtained for various organisms including mammalian cells^10,11^, advances in MS technologies are shifting the focus of phosphoproteomics from a ‘discovery’ mode, towards functionally characterizing the myriad of phosphorylated sites and their kinase-substrate relationships^12,13^. This is a particularly ambitious task considering that < 5% of the more than 100,000 phosphosites currently reported have experimentally-defined cognate kinases, and even fewer are functionally assigned^14^. There is therefore a growing need to further increase the throughput, sensitivity and robustness of MS-based phosphoproteomics workflows to facilitate higher-dimensional experimental designs and to model signaling networks in greater detail.

A pivotal development in proteomics has been the adoption of data-independent acquisition (DIA) methods, which acquire chromatographic elution profiles of peptide fragment ions by cycling through predefined isolation windows encompassing the full precursor mass range^15,16^. While generally producing reproducible datasets, co-isolating multiple precursors within relatively wide isolation windows presents data processing challenges. The most commonly used software suites search for peptides contained in a spectrum library, either built from experimental data, predicted *in silico*, or a combination of both^17–19^. Applications of DIA to phosphoproteomics have demonstrated accurate and reproducible quantification of thousands of phosphosites in time^20,21^ and space^22^ for hundreds of samples per study. We have recently shown that combining trapped ion mobility spectrometry (TIMS) and parallel accumulation – serial fragmentation (PASEF) with DIA (dia-PASEF) enables rapid phosphoproteome measurements without sacrificing depth or sensitivity^23–25^.

As data acquisition speed and robustness improve, scaling sample processing workflows to accommodate more samples becomes a bottleneck. Selective enrichment of phosphorylated peptides from complex biological samples is well-established in proteomics for various affinity chemistries, including immobilized metal cations and metal oxide particles^26,27^. However, state-of-the-art protocols often require specialized equipment for automation and fractionation, or proprietary phosphopeptide enrichment cartridges^8,28–31^. More critically, typical experiments still start with millions of cells per condition to achieve sufficient phosphoproteome depth to cover key regulatory sites, which entails high cost, complexity and significant hands-on processing time. Using smaller sample amounts on the other hand, presents challenges due to lossy sample transfer steps and high volume-to-sample ratios^32–34^.

Here, we introduce µPhos, an efficient, scalable platform for parallel TiO_2_-based phosphopeptide enrichment compatible with cell cultures in 24- to 96-well plates (approximately 20,000 – 100,000 cells). The workflow’s single-pot format minimizes losses from sample transfer steps while reducing the time from living cells to MS injection to one working day. Optimized enrichment and washing conditions enable decreased processing volumes and further enhance sample recovery by reducing peptide-to-surface contact and maintaining sample concentration. In conjunction with highly sensitive dia-PASEF MS, we quantified over 20,000 unique phosphopeptides from 20 µg of starting protein material.

## Materials and methods

### Human cell culture

Human cancer cells (epithelial cervix carcinoma, HeLa) were cultured in RPMI 1640 medium supplemented with 10% fetal bovine serum, 2% penicillin/streptomycin and 5 µl/ml plasmocin in an incubator with 5% CO_2_ at 37 °C. Cells were harvested at ∼80% confluency with 0.25% trypsin/EDTA and collected in 15 ml falcon tubes. Subsequently, cells were washed twice with tris-buffered saline, pelleted by centrifugation for 5 min at 1,000 g and stored at -80 °C until further use.

### Cell lysis for bulk experiments

Cell pellets were lysed in SDC buffer (4% sodium deoxycholate, 100 mM Tris-HCl pH 8.5) and boiled for 7 min at 95 °C before high-energy sonication to shear DNA (Diagenode Bioruptor pico). Protein concentrations were determined via a BCA assay (Thermo Fisher). Protein disulfide bonds were reduced with tris(2-carboxyethyl)phosphine and cysteines were alkylated with 2-chloroacetamide at final concentrations of 10 mM and 40 mM for 5 min at 45 °C.

### µPhos sample processing (protocol style)

1. *Optional* :Cell culture and processing in multi-well plates analogously to the standard protocol described above.
2. Cell lysis aliquots ranging from 1 – 20 µg of starting protein material were diluted to a final volume of 17 µl with SDC lysis buffer in either a 96-deep well or a standard 96-well cell culture plate and sealed with a silicon mat.
3. Cell lysates were sonicated in plates in a water bath sonicator (Elmasonic, S 60/H) for 10 min at room temperature. *Note*: following each sonication or incubation step a quick centrifugation is recommend to ensure lysates are at the base of wells.
4. 2 µl of 10X alkylation/reduction buffer (100 mM and 400 mM of tris(2-carboxyethyl)phosphine and 2-chloracetamide) were added to each well and incubated for 5 min at 45 °C.
5. 1 µl of trypsin/Lys-C mix (final enzyme:protein ratio of approximately 1:100) were added to each well and incubated for at least 2 h at 37 °C.
6. 20 µl of 100% isopropanol were added to each well and mixed for 30 sec at 1,500 rpm in a ThermoMixer.
7. 40 µl of µPhos enrichment buffer (10 mM KH_2_PO_4_, 5 M Glycolic acid in 50% isopropanol) were added to each well and mixed for 30 sec at 1,500 rpm in ThermoMixer.
8. TiO_2_ beads were weighed and suspended in µPhos loading buffer (6% TFA in 80% ACN) at a concentration of 1 mg/µl. 4 µl of the suspension was added to the sample and mixed at 40 °C for 7 min at 1,500 rpm.
9. Beads were pelleted by centrifugation for 1 min at 2,000 g and the supernatant was either stored for proteomics experiments or removed by vacuum aspiration.
10. 200 µl of µPhos washing buffer (5% TFA in 60% isopropanol) was added to beads and incubated for 1 min at 2,500 g in ThermoMixer at room temperature.
11. Beads were pelleted by centrifugation for 1 min at 2,000 g and the supernatant was removed by vacuum aspiration.
12. Steps 10 and 11 were repeated three times.
13. 75 µl of µPhos transfer buffer (0.1% TFA in 60% ACN) were added to each well and the bead suspension was transferred to C_8_ StageTips,
14. Step 13 was repeated.
15. Transfer buffer was removed by centrifugation for 7 min at 1,500 g.
16. Phosphopeptides were eluted with 30 µl of µPhos elution buffer (1% NH_4_OH in 40% ACN) by centrifugation for 4 min at 1,500 g.
17. Step 16 was repeated and the eluates combined.
18. The combined eluates were vacuum centrifuged for 30 min at 45 °C until less than ∼10 µl volume remained.
19. 100 µl of SDB-RPS loading buffer (1% TFA in isopropanol) was added to phosphopeptide solution, loaded on SDB-RPS StageTips and centrifuged for 7 min at 1,000 g.
20. Phosphopeptides were washed with 100 µl of SDB-RPS loading buffer and SDB-RPS washing buffer (0.2% TFA in H_2_O).
21. Phosphopeptides were eluted from StageTips with SDB-RPS elution buffer (1% NH_4_OH in 60% ACN) and vacuum centrifuged to dryness for 40 min at 60 °C.
22. Phosphopeptides were reconstituted in 3 µl MS loading buffer (0.3% TFA, 2% ACN in H_2_O) and stored at -80 °C until LC-MS measurement.

### Liquid chromatography-MS analysis

Purified and desalted peptides were separated via nanoflow reversed-phase liquid chromatography (Bruker Daltonics, nanoElute) within 60 min at flow rate of 0.5 µL/min on a 15 cm x 75 µm column packed with 1.9 µm C_18_ beads (Bruker/PepSep). Mobile phase A was water with 0.1 vol% formic acid and B was ACN with 0.1 vol% formic acid. Peptides eluting from the column were electrosprayed (CaptiveSpray) into a TIMS quadrupole time-of-flight mass spectrometer (Bruker timsTOF HT or timsTOF SCP). We acquired all data in dia-PASEF mode^35^ with an optimized isolation window scheme in the m/z *vs* ion mobility plane for phosphopeptides^24^. The ion mobility range was set from 1/K_0_ = 1.43 to 0.6 Vs cm^-2^ and we used equal ion accumulation time and ramp times in the dual TIMS analyzer of 100 ms each. The collision energy was linearly decreased from 59 eV at 1/K_0_ = 1.4 Vs cm-2 to 20 eV at 1/K_0_ = 0.6 Vs cm^-2^. For all experiments, we calibrated the TIMS elution voltages by known 1/K_0_ values from at least two out of three ions from Agilent ESI LC/MS tuning mix (m/z, 1/K0: 622.0289, 0.9848 Vs cm^-2^; 922.0097, 1.1895 Vs cm^-2^ ; and 1221.9906, 1.3820 Vs cm^-2^).

### Raw data processing

dia-PASEF raw files were processed in Spectronaut v17.0 with spectrum libraries built directly from the DIA experiments (directDIA+). False discovery rates were controlled by a target-decoy approach to ≤1% at precursor and protein levels. We set cysteine carbamidomethylation as a fixed modification and protein N-terminal acetylation, methionine oxidation and serine/threonine/tyrosine phosphorylation as variable modifications. All spectra were matched against the UniProt human reference proteome (accessed 24th April 2022). We activated the PTM localization mode and defined a localization probability score threshold of 0 (‘all phosphopeptides’) or 0.75 (‘Class I phosphopeptides’). Quantification values were filtered by q-value and we defined the ‘Automatic’ normalization mode for cross run normalization. To collapse the Spectronaut output to unique phosphosites, we parsed them with the PeptideCollapse plugin in Perseus^36,37^.

### Bioinformatics analysis

Data analysis and visualization were performed using custom scripts in R (4.0.1) and Python (3.8.8) with packages data.table (1.14.2), dplyr (1.0.7), ggplot2 (3.3.5), tidyR (1.1.14), patchwork (1.1.1), pandas (1.1.5), numpy (1.22.2), plotly (5.4.0), scipy (1.7.3).

## Results and discussion

### Design of a scalable and sensitive phosphoproteomics workflow

The widely used EasyPhos workflow is characterized by streamlined phosphopeptide enrichment in 96-well deep-well plates using TiO_2_ beads, and by the elimination of time-consuming and inefficient fractionation and desalting steps prior to enrichment (**Fig. 1a**). Initially achieved by digesting in a TFE-based buffer directly compatible with downstream enrichment steps^38^, and later by exploiting isopropanol’s ability to prevent precipitation of the normally acid-insoluble detergent SDC under highly acidic conditions^39^, the workflow remains economical and accessible for phosphoproteomics. While the protocol was originally designed for input amounts of >1 mg protein digest, eliminating protein-precipitation steps and optimizing for smaller input materials facilitated phosphopeptide enrichment from hundreds of µg^39^ (**Fig. 1a**). To achieve robust phosphoproteome analysis from tens of thousands of cells, we estimated that it is necessary to increase the entire phosphoproteomics pipeline efficiency from cells to MS by approximately ten-fold (**Fig. 1b**). Similarly to concepts under investigation for single-cell proteomics^40^, we designed µPhos (*i*) to reduce all operation volumes from ∼1 mL to ∼100 µL, (*ii*) to process 96 samples in parallel while minimizing transfer steps and hands-on time, (*iii*) to be compatible with low-input up-front workflows, and (*iv*) to avoid reliance on specialized equipment or reagents, maximizing accessibility of the workflow.

**Figure 1.**
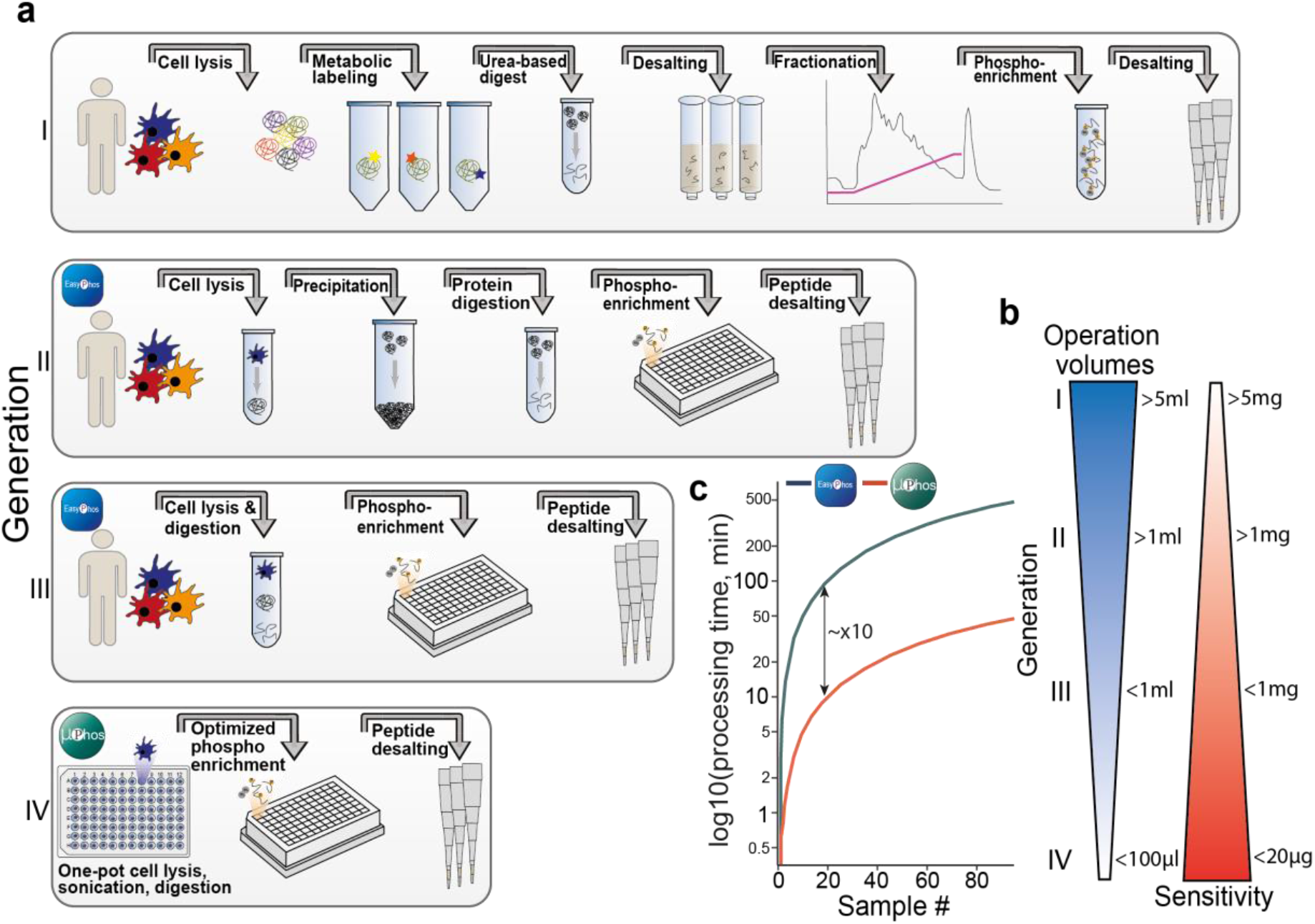
Design of the µPhos platform. **a** Schematic overview of the development of TiO_2_-based protocols for MS-based phosphoproteomics. **b** Comparison of typical protein starting amounts and phosphopeptide enrichment volumes for the protocols in a. **c** Estimation of the cumulative time-per-sample for pre-digestion processing steps calculated from the average of 96 samples.

We first examined the timings of a typical phosphoproteomics experiment (**Suppl.Fig. 1**). In our hands, processing 96 samples according to the EasyPhos protocol (excluding digestion time) required on average about 5.2 min per sample (**Fig. 1c**). A substantial fraction of this time was consumed by upstream sample preparation steps, including cell harvesting, lysis, and the routinely performed high-energy sonication step for DNA shearing. This step, unless using specialized instrumentation, cannot be readily parallelized for >12 samples, presenting a major bottleneck for up-scaling. However, when working with lower input materials in smaller volumes, we found water-bath sonication could replace this step without impacting sample quality. We also found it possible to omit protein concentration determination and several transfer steps. As a result, we reduced the average time per sample to only 0.5 min in the µPhos protocol. Although insignificant for small sample numbers, this substantially improves feasibility of preparing many samples in large-scale studies. Overall, these improvements should enhance scalability, reproducibility and data quality.

### Optimized phosphopeptide enrichment in small volumes

High-sensitivity proteomics workflows often aim to reduce non-specific adsorption to plastic surfaces by working in minimal volumes. To assess this in the context of phosphoproteomics, we progressively decreased the total volume at the enrichment step and assessed the number of identified peptides with dia-PASEF (**Fig. 2**). In the high-sensitivity EasyPhos protocol^39^, lysis and digestion buffer, isopropanol and enrichment buffer contribute to ∼800 µL during enrichment. Analyzing 20 µg HeLa cell lysates and progressively reducing all volumes proportionally to 80 µL led to a substantial increase in the median fragment ion current, plateauing at 40 µL (**Fig. 2a**). Accordingly, we observed an almost linear increase in the number of identified peptides (**Fig. 2b**). However, the relative proportion of non-phosphorylated peptides also increased more strongly than the number of phosphorylated peptides, resulting in a lower enrichment selectivity as enrichment volumes decreased. Further investigation of unmodified (**Fig. 2c**) and phosphorylated (**Fig. 2d**) peptide intensities from our MS measurements showed that the median intensity of unmodified peptides increased almost 30-fold from 800 to 40 µL enrichment volume, while the intensity of phosphorylated peptides increased only 4-fold in the same range. Thus, although our data supported the concept of increasing sensitivity by minimizing enrichment volumes, this came at the expense of reduced enrichment selectivity. Bearing this in mind and to ensure compatibility with standard 96-well plates, we chose 80 µL as the working volume for subsequent experiments.

**Figure 2.**
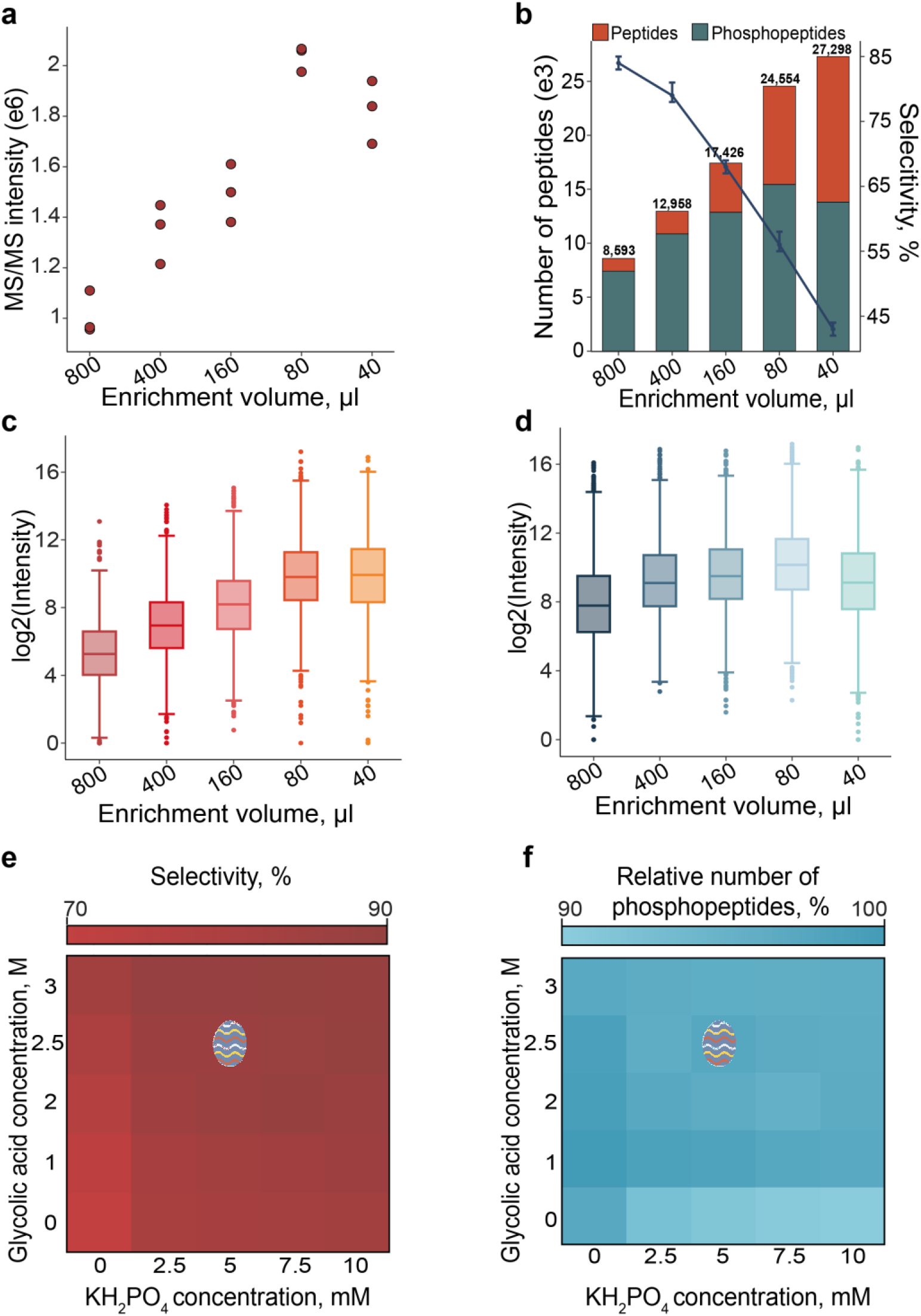
Optimization of the phosphopeptide enrichment. **a** Median raw fragment ion intensity in three replicate injections as a function of enrichment volume. **b** Number of identified unmodified and phosphopeptides for decreasing enrichment volumes starting from 20 µg HeLa cell lysates. The line plot shows the selectivity of the phosphopeptide enrichment in percent. **c** Logarithmized intensity of unmodified peptides for phosphopeptide enrichments with decreasing volume. **d** Same as c, but for phosphopeptides. **e** Selectivity matrix for glycolic acid and KH_2_PO_4_ as selectivity agents. **f** Same as e, but showing the relative number of identified phosphopeptides.

It is well established that the binding equilibrium of phosphorylated peptides to TiO_2_ can be influenced by various modifiers^41,42^. The most recent EasyPhos protocol employed 1 mM monopotassium phosphate (KH_2_PO_4_) to compete with non-phosphopeptide binding. We found that under the new enrichment conditions (20 µg starting material and 80 µL enrichment volume), increasing the KH_2_PO_4_ concentration proportionally up to 10 mM improved the relative fraction of phosphopeptides versus non-phosphorylated peptides (**Suppl. Fig. 2a**). However, this also reduced the total number of identified phosphopeptides, presumably due to competition between the KH_2_PO_4_ and phosphorylated peptides. We therefore explored hydrophilic organic acids such as glycolic or lactic acid, which have been commonly used as MS-compatible modifiers^41,43^. Screening various concentrations and combinations suggested a mixture of 1 M glycolic, 1 M lactic acid and 5 mM KH_2_PO_4_ as a promising starting point for further optimization (**Suppl. Fig. 2b**). To simplify the protocol, we omitted lactic acid and focused on a binary combination of glycolic acid and KH_2_PO_4_. Balancing selectivity (**Fig. 2e**) with the number of identified phosphopeptides (**Fig. 2f**), we independently varied the final concentrations of glycolic acid and KH_2_PO_4_ from 0 to 3 M and 0 to 10 mM, respectively. Interestingly, when used as single agents, both compounds performed sub-optimally. However, combining at least 2 M glycolic acid and 5 mM KH_2_PO_4_ resulted in a robust plateau with selectivity >85% and phosphopeptide identifications varying less than 5%. We thus selected 2.5 M glycolic acid and 5 mM KH_2_PO_4_ as our default enrichment modifiers. As a result, µPhos enables phosphopeptide enrichment in 10-fold lower volumes without compromising selectivity, while maintaining isopropanol concentration high enough to prevent SDC precipitation, allowing all steps to be performed in a single reactor.

### Reproducible one-day protocol for phosphoproteomics

Having established conditions for phosphopeptide enrichment in a more scalable format, we aimed to perform the complete phosphoproteomics workflow in one day. Traditionally, proteolytic digestion is performed overnight, but shorter digestion times of a few hours are frequently explored for studies involving very small sample amounts such as single cells^44^ or for clinical proteomics^45^. Until now, there has been little incentive to apply this approach to phosphoproteomics due to the relatively large input amounts and time-consuming upstream sample preparation. We therefore digested 10 µg of HeLa cell lysates for 2, 4 and 8 hours (mimicking overnight digestion), using three different enzyme:protein ratios ranging from 1:50 to 1:200 for both LysC and trypsin. All conditions resulted in a similar number of phosphopeptides with no clear trend along either axis, but the fraction of phosphorylated peptides with more than one missed tryptic cleavage site decreased slightly with increasing duration and enzyme concentration (**Fig. 3a**). We concluded that digestion for 4 h at an enzyme:protein ratio of 1:50 yields an adequate digestion efficiency (<8% peptides with > 1 internal lysine or arginine), while still allowing the entire protocol to be completed within one working day. The physicochemical properties of identified peptides were virtually identical even between the 2 h and 8 h digestions, further supporting this point (**Fig. 3b**). Additionally, all digestion conditions yielded similar quantitative results, with median coefficients of variation around 25% in three replicates (**Fig. 3c**).

**Figure 3.**
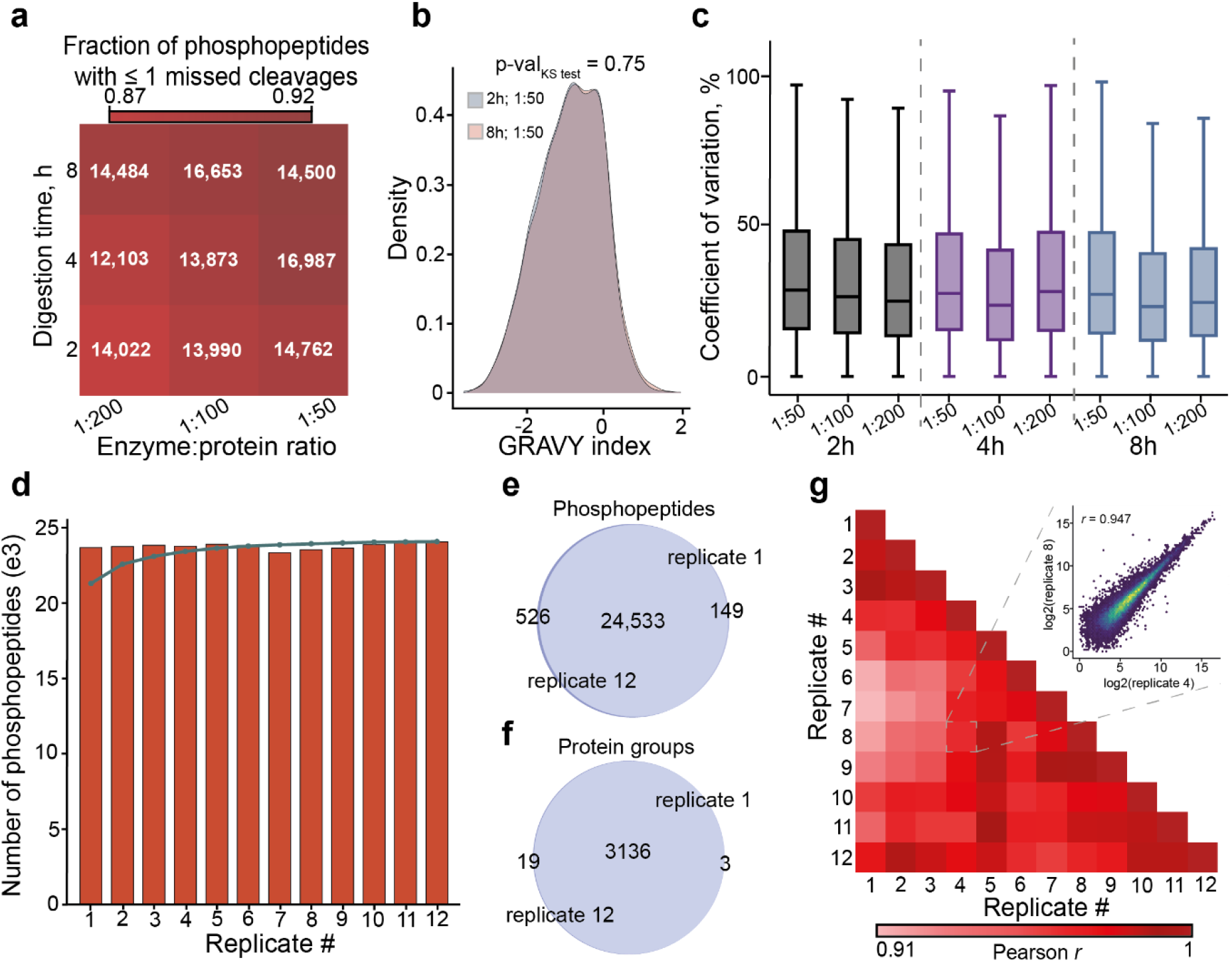
Robustness and scalability of the µPhos platform. **a** Digestion efficiency and number of identified phosphopeptides in a matrix of incubation time and enzyme:protein ratio for starting amounts of 10 µg HeLa lysate. **b** Overlay of the GRAVY hydrophobicity index of detected phosphopeptides after either 2h or 8h digestion. **c** Precision of label-free phosphopeptide quantification in workflow replicates (n=3) for the conditions in a. Outliers not shown. **d** Number of identified phosphopeptides in twelve workflow replicates starting from 10 µg HeLa lysate. The blue line indicates the cumulative number. **e** Overlapping phosphopeptide identifications in the first and last replicate from d. **f** Same as e, but phosphoproteins. **g** Pairwise Pearson-correlation matrix for phosphopeptides quantified in twelve workflow replicates starting from 10 µg HeLa lysate and digesting for 4 h with a 1:50 protein:enzyme ratio. The inset shows a scatter plot for a pair of replicates with a correlation coefficient close to the median.

Considering large-scale applications, we next tested the reproducibility of µPhos for *n* > 3 samples. We prepared a single batch of 12 workflow replicates from aliquots of one HeLa cell lysate with the new digestion and enrichment conditions. Similar to the previous results, we observed a highly consistent performance, identifying 23,000 ± 150 phosphopeptides per replicate (**Fig. 3d**). Out of these, 12,477 phosphosites were localized with a probability score >0.75. The replicates exhibited a very high overlap of both peptide and phosphoprotein identifications (**Fig. 3e-f**), resulting in an almost complete quantification matrix with only 2.5% missing values. Comparing phosphopeptide intensities pairwise across replicates, we found a median Pearson correlation coefficient of 0.93 (**Fig. 3g**) and a median coefficient of variation of 19%. For reference, the latter is just five percentage points higher than what we recently found when assessing only the technical variability of label-free quantification in 7-min dia-PASEF runs with ten replicate injections of pooled phosphopeptides.

### Application to limited sample amounts

These results indicate that µPhos is compatible with significantly lower input amounts than those typically used in phosphoproteomics studies. We performed a dilution series experiment, ranging from 20 to 1 µg aliquots of a HeLa cell lysate, to determine the platform’s sensitivity limits. We used dia-PASEF on a timsTOF HT mass spectrometer and a timsTOF SCP system, specifically designed for ultra-high sensitivity applications^46^. The SCP’s brighter ion source led to increased ion current and phosphopeptide intensity (**Fig. 4a**). For 20 µg starting material, the difference in peptide intensity between the systems was 2.85-fold, which decreased to 2.0- and 1.9-fold at 10 and 5 µg, and was no longer evident at 1 µg. This translated into 23,100 and 23,800 unique phosphopeptide sequences for the 20 µg sample on the HT and SCP systems respectively, with over 12,000 localized sites (localization probability >0.75) for both (**Fig. 4b**). With half the input amount, over 15,000 phosphopeptides and 8,000 class 1 sites were still quantified. At 5 µg, the SCP instrument outperformed the timsTOF HT by 35% phosphopeptides and 30% class 1 sites. However, we observed diminishing returns for the more sensitive mass spectrometer at 1 µg, suggesting sample processing may again be a limiting factor with very low input material.

**Figure 4.**
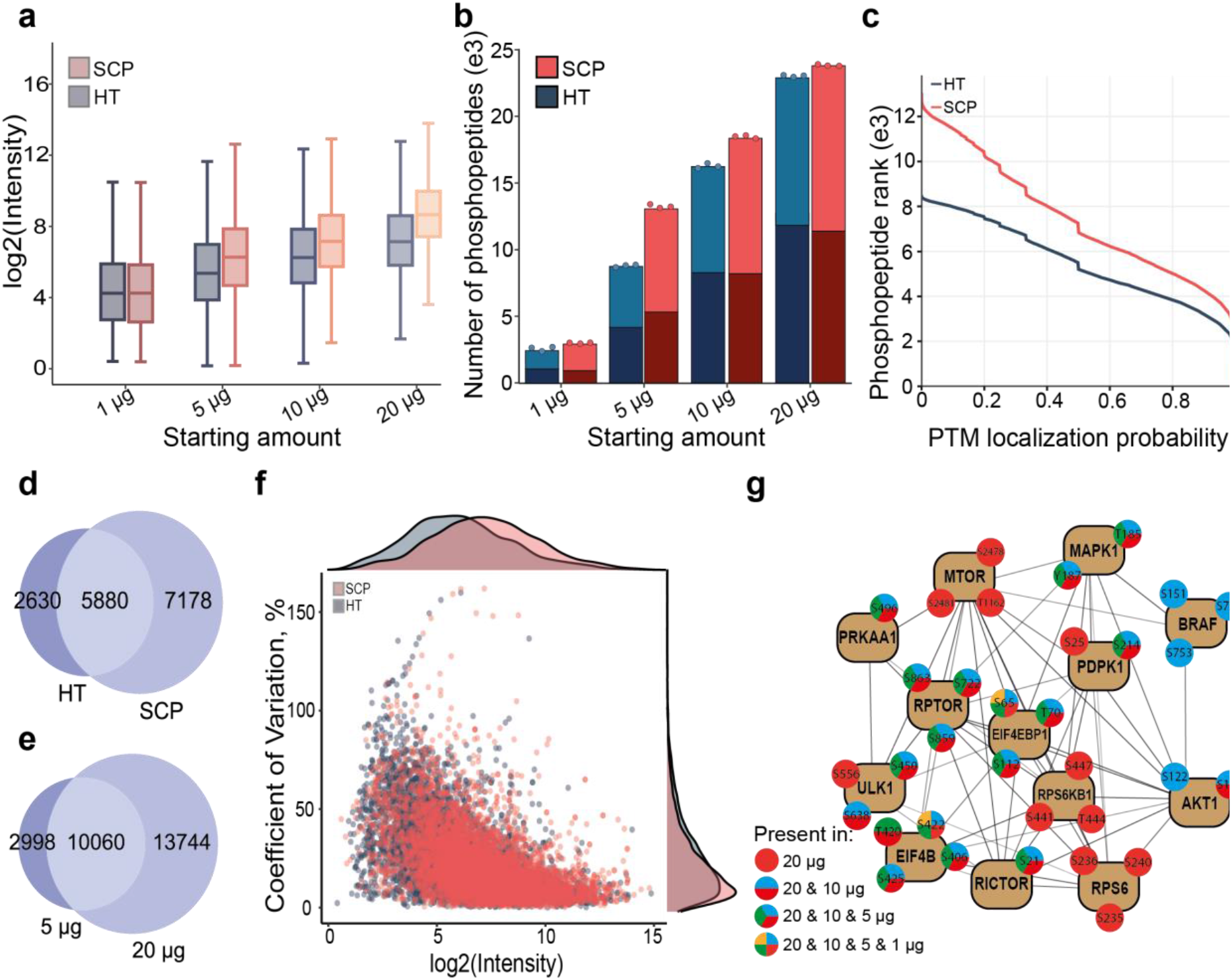
Sensitive enrichment of phosphopeptides from low-µg starting amounts. **a** Logarithmized phosphosite intensities acquired on timsTOF HT and timsTOF SCP. **b** Number of identified phosphopeptides and Class I phosphosites (localization score >0.75, darker color) as a function of protein starting amount. **c** Number of phosphopeptides as a function of the Spectronaut site localization probability score. **d** Overlapping phosphopeptide identifications from 5 µg of starting amount acquired on timsTOF HT and timsTOF SCP. **e** Overlapping phosphopeptide identifications from 5 µg and 20 µg starting amount acquired on timsTOF SCP. **f** Coefficients of variation for the quantified overlapping phosphopeptides from **d** as a function of their mean intensity. Density plots are projected on the respective axis. **g** Scheme of protein phosphorylation sites associated with the KEGG term ‘mTOR signaling pathway’ and quantified in µPhos experiments starting from 1, 5, 10 and 20 µg lysate.

Closer examination of the MS data at 5 µg showed that the SCP instrument’s higher signal resulted in more phosphopeptides identified at any localization score cut-off (**Fig. 4c**). That is, even with a stringent localization probability cut-off of >0.99, we could identify ∼3,000 phosphosites. Approximately two-thirds of all unique phosphopeptide sequences identified with the HT overlapped with the SCP instrument, which identified ∼7,000 additional unique sequences from the same starting amount (**Fig. 4d**). As expected, phosphopeptides identified from 5 µg were largely a subset of the 20 µg data (**Fig. 4e**). Further, our data revealed a higher quantitative reproducibility for the timsTOF SCP at 5 µg, particularly for low-abundance phosphopeptides (**Fig 4f**).

To demonstrate the biological information that can be obtained from low input amounts, we focused on the well-studied mammalian target of rapamycin (mTOR) signaling pathway, a key regulator of cell growth and metabolism^47^. We filtered proteins associated with the KEGG term ‘mTOR signaling pathway’ from Class I sites in the timsTOF SCP dataset. **Figure 4g** shows curated phosphosites in PhosphoSitePlus of core members of the mTOR complexes and upstream or downstream kinases. Of these, we quantified 78 phosphosites from 20 µg starting material, which reduced to ∼40 at 10 and 5 µg. Only 14 sites were quantifiable from 1 µg input material. Assuming roughly 250 pg protein amount per HeLa cell^48^, we concluded that the combination of µPhos with highly sensitivity TIMS MS effectively quantifies biologically relevant phosphosites from ≥20,000 cells (∼5 µg).

## Discussion

The transition of MS-based phosphoproteomics from mapping phosphorylation sites to their functional characterization requires increased throughput, reduced input amounts and streamlined sample processing. To meet these demands, we developed µPhos, a robust and accessible protocol optimized for protein yields from 20,000 to 100,000 cells. Its key advantage is the miniaturization of upstream cell lysis and proteolytic digestion, enabling phosphopeptide enrichment from crude digests in < 100 µL total volume without intermediate steps. This was achieved by optimizing enrichment conditions to minimize non-specific binding of unphosphorylated peptides at high bead concentrations. Under these conditions, we also found that the duration of tryptic digestion could be reduced from overnight to just 4 hours. Together with eliminating other time-consuming steps, this workflow now enables processing of 96 samples within one working day at a per-sample cost of ∼$10 for consumables and reagents.

Minimizing sample handling steps should also reduce variability within and between experiments. We found that technical reproducibility, as estimated by phosphopeptide coefficients of variation, is comparable to other state-of-the-art workflows for conventional starting amounts^32,49^. Further improvements to throughput can be achieved by automation, for example with magnetic beads^30^. However, such adaptations require increasingly specialized equipment and proprietary reagents. They also require careful re-evaluation of enrichment conditions, as different bead chemistries and physiochemical properties strongly influence phosphopeptide enrichment performance even under standardized conditions^50^. Nonetheless, while we optimized the µPhos protocol using standard TiO_2_ material, exploring the performance of the workflow using alternative chemistries specifically in high-sensitivity applications with low enrichment volumes may be rewarding.

In addition to facilitating parallel processing of numerous samples, reducing sample preparation volumes and minimizing exposure to surfaces generally alleviates adsorptive peptide losses, thereby increasing sample recovery. We observed a 4-fold increase in raw MS intensity compared to the previous generation protocol. However, single-cell proteomics workflows routinely process samples in 5 µL or less^51,52^, suggesting further sensitivity gains may be achievable through continued miniaturization. One practical limitation is the need for sufficient volumes to collect all cells while ensuring efficient lysis and mixing during enrichment. Nonetheless, the sensitivity of our label-free analysis of low µg inputs from an unperturbed cell line using a timsTOF mass spectrometer compares favorably with other approaches tailored to minimal sample amounts^32–34,49^, and outperforms them with > 5 µg starting material. Thus, while robust phosphoproteomics from 1 µg or less remains challenging, µPhos shifts the working range from ∼200 µg to ∼20 µg for most practical applications.

Advances in bioinformatics such as the integration of neural networks have made the direct analysis of DIA data without the need for experimental libraries a preferred choice in proteomics studies^18,29,53,54,55^. Given the combinatorial challenges in phosphoproteomics, library-based approaches may still provide superior coverage for small numbers of samples, but we expect this advantage, if any, to diminish as the number of samples increases. At present, label-free quantification appears most suitable for large-scale phosphoproteomics studies, as it can simply scale to compare as many samples as required^56^. It remains to be explored whether recent multiplexing strategies can also be combined with µPhos, which may be particularly beneficial for sub-µg input amounts^32,52,57^; however, in principle the workflow should be adaptable to either isobaric or non-isobaric labeling.

We conclude that µPhos is well suited for large-scale phosphoproteomics studies and anticipate a wide range of applications providing new insights into cellular signaling networks, including their responses to internal or external perturbations, or as a function of their cellular and spatial context. The ability to process 96 samples within one working day, combined with high reproducibility and phosphoproteome coverage from minimal starting material will likely contribute to the acceleration of phosphoproteomics research, opening new avenues for studying signal transduction.

## Acknowledgements

This work was partially supported by the Federal Ministry of Education and Research and the Thuringian Ministry for Economic Affairs, Science and a Digital Society through the Joint Federal Government-Länder Tenure-Track Programme, by the Free state of Thuringia and the European Union via the “Innovationszentrum für Thüringer Medizin technik-Lösungen” (ThIMEDOP; #2018 IZN 002), by the Center for Interdisciplinary Clinical Research (IZKF Jena),by the German Research Foundation through the Research Training Group ‘ProMoAge’ (RTG 2155), by the National Health and Medical Research Foundation (NHMRC; grant #2011083), and the Stafford Fox Medical Research Foundation. We acknowledge L. Reiter, T. Gandhi and O. Bernhardt (BiognoSYS AG) for technical support with Spectronaut and T. Müller, M. Lubeck and O. Räther (Bruker) for technical support and discussions. We are grateful to our colleagues at the Jena University Hospital for fruitful discussions and technical support, in particular C. Tschernjawski.

## Supplementary Information

**Supplementary Figure 1.**
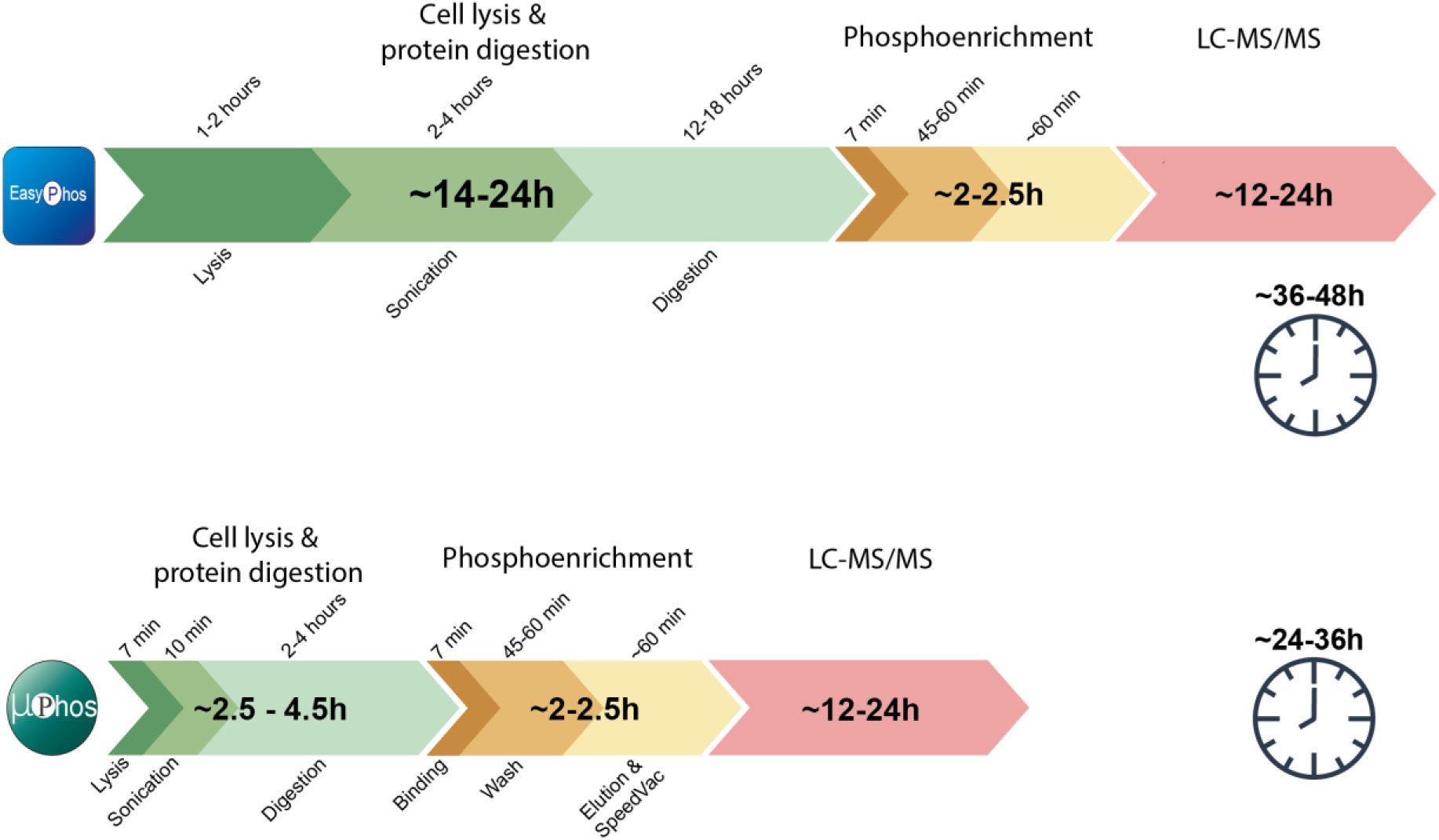
Typical processing times for EasyPhos and µPhos protocols.

**Supplementary Figure 2.**
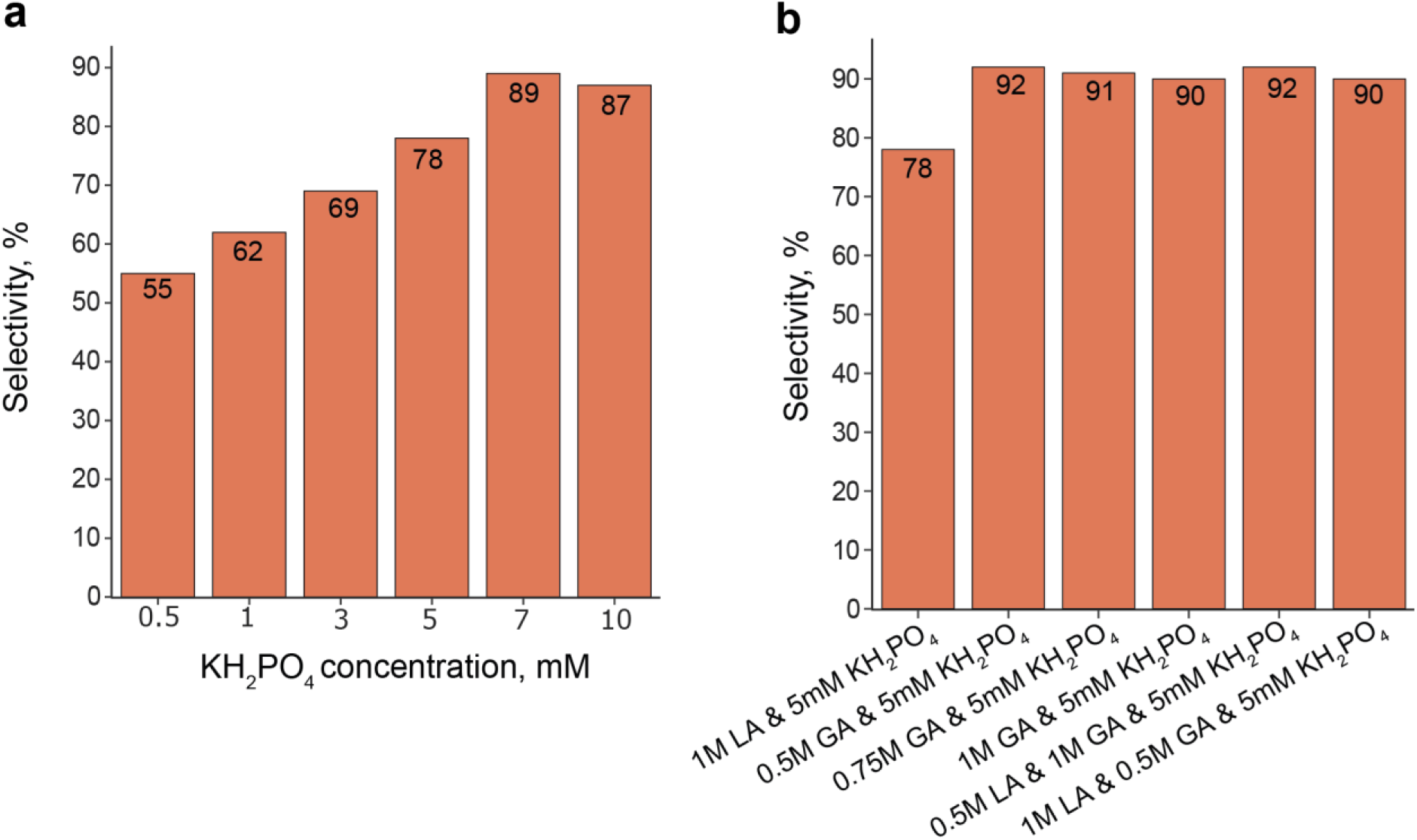
**a** Selectivity of phosphoenrichment as a function of increased concentration of monopotassium phosphate. **b** Same as a but for combination of selectivity agents

## Notes

### Competing Interest Statement

The authors have declared no competing interest.

